# The short- and long-range RNA-RNA Interactome of SARS-CoV-2

**DOI:** 10.1101/2020.07.19.211110

**Authors:** Omer Ziv, Jonathan Price, Lyudmila Shalamova, Tsveta Kamenova, Ian Goodfellow, Friedemann Weber, Eric A. Miska

## Abstract

The *Coronaviridae* is a family of positive-strand RNA viruses that includes SARS-CoV-2, the etiologic agent of the COVID-19 pandemic. Bearing the largest single-stranded RNA genomes in nature, coronaviruses are critically dependent on long-distance RNA-RNA interactions to regulate the viral transcription and replication pathways. Here we experimentally mapped the *in vivo* RNA-RNA interactome of the full-length SARS-CoV-2 genome and subgenomic mRNAs. We uncovered a network of RNA-RNA interactions spanning tens of thousands of nucleotides. These interactions reveal that the viral genome and subgenomes adopt alternative topologies inside cells, and engage in different interactions with host RNAs. Notably, we discovered a long-range RNA-RNA interaction - the FSE-arch - that encircles the programmed ribosomal frameshifting element. The FSE-arch is conserved in the related MERS-CoV and is under purifying selection. Our findings illuminate RNA structure based mechanisms governing replication, discontinuous transcription, and translation of coronaviruses, and will aid future efforts to develop antiviral strategies.

## INTRODUCTION

RNA viruses comprise the dominant component of the eukaryotic virome (Dolja and Koonin, 2018). Their error-prone genome replication mode allows them to rapidly evolve new variants and to jump from animals to humans (Woolhouse and Gaunt, 2007), thus presenting a high epidemic and pandemic threat. Several members of the betacoronavirus genus (family *Coronaviridae*), namely the <u>S</u>evere <u>A</u>cute <u>R</u>espiratory <u>S</u>yndrome <u>co</u>rona<u>v</u>irus (SARS-CoV), the <u>M</u>iddle <u>E</u>ast <u>R</u>espiratory <u>S</u>yndrome <u>co</u>rona<u>v</u>irus (MERS-CoV), as well as the <u>S</u>evere <u>A</u>cute <u>R</u>espiratory <u>S</u>yndrome <u>co</u>rona<u>v</u>irus <u>2</u> (SARS-CoV-2) are of special concern. SARS-CoV-2, the causative agent of <u>Co</u>rona<u>vi</u>rus <u>D</u>isease 20<u>19</u> (COVID-19), has spread to date to nearly every country in the world, resulting in millions of infections, over a million of deaths, and a massive global economic impact (McKibbin and Fernando, 2020). Even though worldwide efforts and resources are redirected to overcome the COVID-19 pandemic, at present, there are no approved vaccines or antiviral medicines. This illustrates the urgent need for deciphering in-depth the molecular biology of coronaviruses, especially SARS-CoV-2.

Coronaviruses have evolved the largest known single-stranded RNA genome in nature. Regulation of their mRNA transcription and translation is facilitated by cis-acting structures that interact with each other, with viral proteins, and with host machineries (Madhugiri et al., 2016). mRNA transcription in coronaviruses involves a process whereby so-called <u>s</u>ub<u>g</u>enomic <u>mRNAs</u> (sgmRNAs) are produced through discontinuous <u>g</u>enomic <u>RNA</u> (gRNA) template utilization, which is in contrast to replication of the full-length genome (Sawicki et al., 2007). This discontinuous transcription is mediated by the <u>T</u>ranscription <u>R</u>egulating <u>S</u>equence-leader (TRS-L) at the 5′ end of the genome, and the <u>T</u>ranscription <u>R</u>egulating <u>S</u>equence-<u>b</u>ody (TRS-B) at the 5′ ends of each ORF. Template switching between these RNA sequence elements results in a set of 5′ and 3′ co-terminal, “nested” sgmRNAs of different sizes on which the 5′ proximal ORFs are translated into nonstructural or structural viral proteins (Moreno et al., 2008; Sola et al., 2015). The mechanisms underlying discontinuous transcription and genome replication have not been fully worked out, however long-distance RNA-RNA interactions along the viral genome have been proposed as key regulators (Mateos-Gómez et al., 2011; Mateos-Gomez et al., 2013; Moreno et al., 2008; Sola et al., 2015).

On the full-length gRNA itself, two partially overlapping open reading frames (ORF1a and ORF1b) are translated from the same start codon at the 5′ end, resulting in the polyproteins pp1a and pp1ab. Translation of the longer product pp1ab is made possible by a hairpin-type pseudoknot RNA structure known as the frame<u>s</u>hifting <u>e</u>lement (FSE) which regulates a programmed −1 ribosomal frameshifting that overrides with about 50% efficiency the stop codon of ORF1a (Kelly et al., 2020; Namy et al., 2006). Previous studies applied RNA structure probing techniques using selective 2′-hydroxyl acylation analyzed by primer extension (SHAPE) and DMS, as well as nuclear magnetic resonance (NMR) to effectively identify conserved cis-acting RNA structures regulating the life cycle of coronaviruses. However, when it comes to identifying long-distance base-pairing between distal nucleotides, these methods fall short. Therefore, the long-range RNA-RNA interactome of coronaviruses has never been mapped in full. Deciphering how the various structural elements along the coronavirus gRNA and sgmRNA are folded and brought together in time and space is vital for understanding, dissecting and manipulating viral replication, discontinuous transcription, and translation regulation.

We recently developed <u>C</u>rosslinking <u>O</u>f <u>M</u>atched <u>R</u>NAs <u>A</u>nd <u>De</u>ep <u>S</u>equencing (COMRADES) for in-depth RNA conformation capture in living cells (Ziv et al., 2018). COMRADES is derived from a class of methods that combine psoralen crosslinking of base paired RNA and deep sequencing (Aw et al., 2016; Lu et al., 2016; Sharma et al., 2016). COMRADES utilizes a clickable psoralen derivative to specifically crosslink paired nucleotides, and high throughput sequencing to retrieve their positions (Figure 1). Following *in vivo* crosslinking, the viral RNA is selectively captured, fragmented and subjected to a click-chemistry reaction to add a biotin tag to crosslinked fragments. Crosslinked RNA duplexes are then selectively captured using streptavidin affinity purification. Half of the resulting RNA is proximity ligated, following reversal of the crosslink to create chimeric RNA templates for high throughput sequencing. The other half is used as a control, in which reversal of the crosslink precedes the proximity ligation, and accurately represents the background level of non-specific ligation. The coupling of two biotin-streptavidin mediated enrichment steps, first of viral RNA, and second of crosslinked RNA duplexes provides high structural depth for identification of both long- and short-lived conformations. COMRADES can therefore measure *(i)* the structural diversity of alternative RNA conformations that co-exist inside cells; *(ii)* short-distance, as well as long-distance (over tens of thousands of nucleotides) base-pairing within the same RNA molecule; and *(iii)* base-pairing between different RNA molecules, such as those of host and viral origin (Kudla et al., 2020; Ziv et al., 2018).

**Figure 1.**
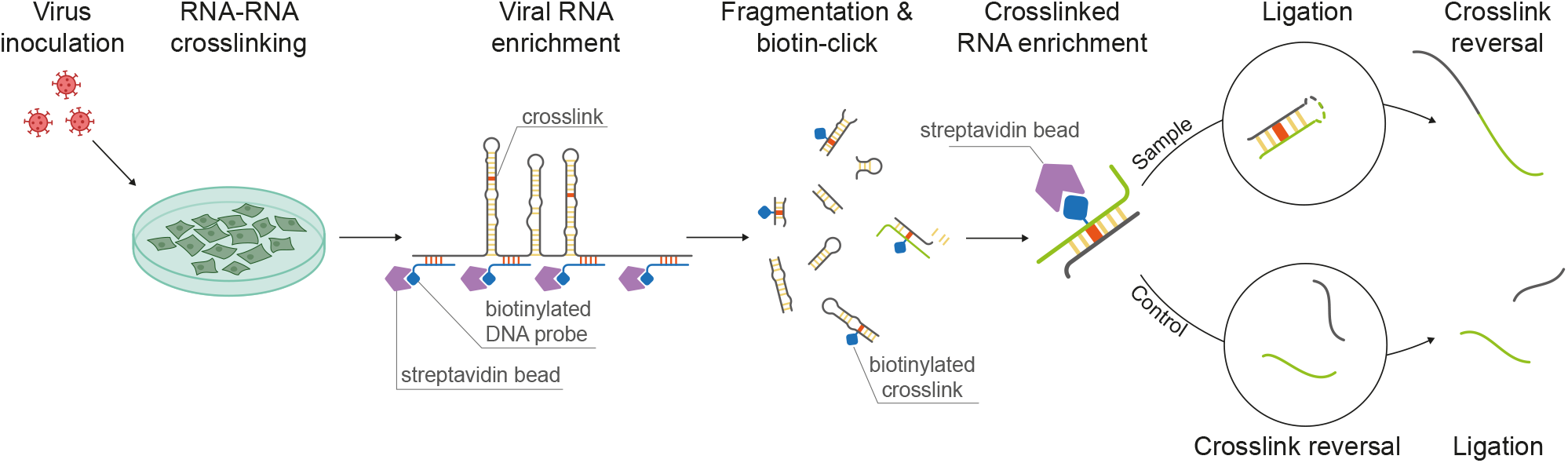
The COMRADES method. Virus inoculated cells are crosslinked using clickable psoralen. Viral RNA is pulled down from the cell lysate using an array of biotinylated DNA probes, following digestion of the DNA probes and fragmentation of the RNA. Biotin is attached to crosslinked RNA duplexes via click chemistry, enabling pulling down crosslinked RNA using StreptAvidin beads. Half of the RNA duplexes are proximity ligated, following reversal of the crosslinking to enable sequencing. The other half serves as a control, in which crosslink reversal proceeds the proximity ligation. See Figure S1 for numbers and percentages of chimeric reads.

Here we apply COMRADES to study the structural diversity of the SARS-CoV-2 gRNA and sgmRNA inside cells. We discover networks of short- and long-range RNA-RNA interactions spanning the entirety of SARS-CoV-2 gRNA and sgmRNA. We reveal site-specific interactions with the host transcriptome. Finally, we uncover a conserved long-range structure encompassing the programmed ribosomal frameshifting element.

## RESULTS AND DISCUSSION

### The SARS-CoV-2 genome and sgmRNA adopt alternative co-existing topologies that involve long-distance base-pairing

Inside the host, the gRNA of SARS-CoV-2 is transcribed into sgmRNA (Figure 2A). To compare the structure of both types of RNA, we applied the COMRADES method and set up a dual enrichment strategy to analyse the positive sense gRNA and positive sense sgmRNA separately (Figure 2B). Briefly, we selectively pulled down the full-length positive sense SARS-CoV-2 genome from *in vivo* crosslinked, SARS-CoV-2 inoculated Vero E6/TMPRSS2 cells (Matsuyama et al., 2020), using a tiling array of antisense probes for ORF1a/b, which resulted in a highly enriched gRNA fraction (Figure 2C). The full-length positive sense sgmRNA was subsequently enriched from the gRNA-depleted supernatant of the first pulldown, using a second tiling array of antisense probes to the region downstream of ORF1a/b (Figure 2B). This dual enrichment strategy resulted in a high degree of separation between the gRNA and the sgmRNA (Figure 2C). COMRADES provided >6 million non-redundant chimeric reads, which was sufficient to generate high-resolution maps for both the gRNA and sgmRNA with a high signal to noise ratio (Figure S1), and high reproducibility between independent biological replicates (r = 0.92, p value <2.2e-16, Figure 2D,E). Our structural data covered >99.99% of the coronavirus gRNA and the sgmRNA (Figure 2C), and represents the base-pairing nature of SARS-CoV-2 gRNA and sgmRNA inside cells.

**Figure 2.**
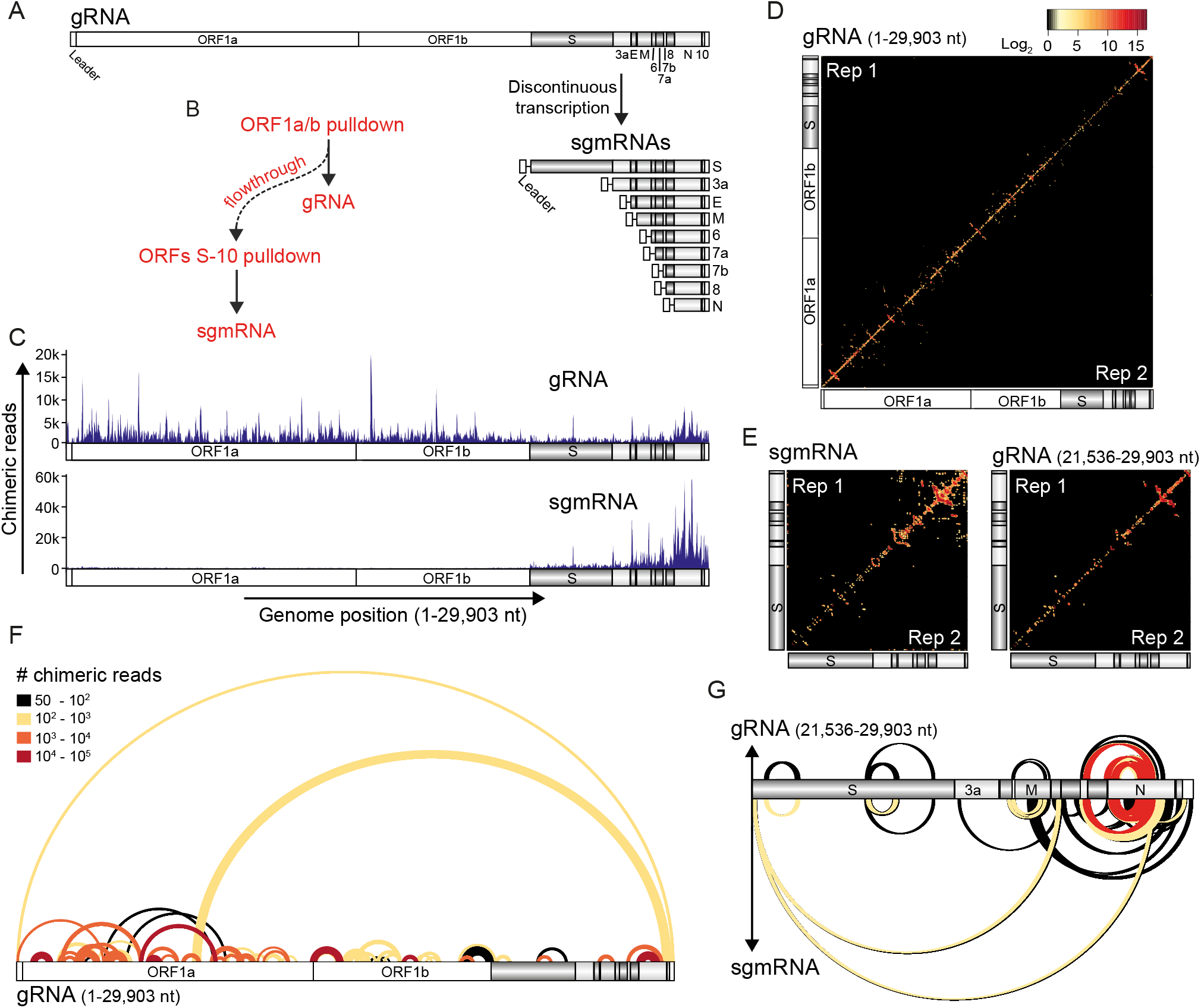
The *in vivo* base-pairing of the SARS-CoV-2 gRNA and sgmRNA. (A) The organisation of the SARS-CoV-2 gRNA and sgmRNA. (B) Dual-enrichment strategy for separation of SARS-CoV-2 gRNA and sgmRNA from inside cells. (C) Coverage of structural data (chimeric reads) for the gRNA and sgmRNA samples aligned to the gRNA coordinates. Each chimeric-read originated from *in vivo* crosslinking of base-paired RNA. Chimeric reads aligned to the Leader in the sgmRNA samples are not shown due to the use of gRNA coordinates. (D) Heat map of RNA-RNA interactions along the SARS-CoV-2 gRNA. Signal represents base-pairing between the genomic coordinates on the *x* and *y* axes. Top left and bottom right represent two independent biological replicates. Colour code corresponds to the number of non-redundant chimeric reads supporting each interaction. (E) Left panel: Heatmap of RNA-RNA interactions along the ORF S sgmRNA. Right panel: Zoom-in of the gRNA region from (D) corresponding to ORF S sgmRNA coordinates. Colour code as in (D). Top left and bottom right represent two independent biological replicates. (F) Arch plot representation of long-range RNA-RNA interactions along the SARS-CoV-2 gRNA. Interactions that span at least 500 nt are shown. Colours represent the number of non-redundant chimeric reads supporting each arch. (G) Arch plots representation of long-range RNA-RNA interactions along ORF S sgmRNA (bottom) and the gRNA region corresponding to the S sgmRNA coordinates (top). Interactions that span at least 500 nt are shown. Colours as in (F).

Available models for the RNA structure of SARS-CoV-2 and related viruses are largely confined to short-distance base-pairing which result in local folding of important cis-acting elements (Andrews et al., 2020; Huston et al., 2020; Kelly et al., 2020; Lan et al., 2020; Manfredonia et al., 2020; Ryder, 2020; Sanders et al., 2020; Sun et al., 2020). However, long-distance base-pairing between distal RNA elements are equally essential for many RNA viruses (Huber et al., 2019), including coronaviruses (Mateos-Gómez et al., 2011; Mateos-Gomez et al., 2013; Moreno et al., 2008; Sola et al., 2015). The ability of COMRADES to capture RNA base-pairing regardless of the distance between the interacting bases enabled us to confirm *in vivo* the structure of nearly all previously characterised cis-acting elements (with one exception, discussed below) and to discover long-distance RNA-RNA interactions as they occur inside cells. Indeed, we observed a high prevalence of long-range RNA base-pairing along the SARS-CoV-2 genome, with ORF1a demonstrating more long-range connectivity than any other ORF (Figure 2F). Most of the base-pairing is confined to a single ORF, however, some interactions cross ORF boundaries. For example, ORF1a base-pairs with ORF1b, as well as with the 5′ and 3′ untranslated regions (UTRs) (Figure 2F). We additionally discovered long-distance interactions unique to the sgmRNA (Figure 2G). Previous models of the SARS-CoV-2 and related viruses mainly analysed structural population averages, i.e. assuming that all copies of the genome and sgmRNA have a single static conformation. Yet, the complex life cycle of viral RNA genomes, i.e. their engagement with multiple cellular and viral machineries such as the ones for replication, transcription, and translation, suggests a dynamic RNA structure, as we and others have reported for Zika virus (Huber et al., 2019; Li et al., 2018; Ziv et al., 2018) and for HIV-1 (Tomezsko et al., 2020). Our structural analysis of SARS-CoV-2 reveals a high level of structural dynamics whereby alternative high-order conformations, some of which involve long-distance base-pairing, co-exist *in vivo* (Figures 3 and S2A, Table S1). For example, nucleotides 5,660-5,680 in ORF1a interact with three alternative distal regions: 3.6 kb upstream, 3.4 kb downstream, and 2 kb upstream (Figure 3, arches 4, 5 and 8 respectively), and the 5′ UTR interacts with ORF1a as well as with the 3′ UTR (Figure 3, arches 2 and 3 respectively). In contrast, we find that ORF N sgmRNA is held in a single dominant conformation where the leader sequence interacts exclusively with a region 0.8 kb downstream (Figure S2B). In summary, we discover the co-existence of alternative SARS-CoV-2 gRNA and sgmRNA topologies, held by long-range base-pairing between regions tens of thousands of nucleotides apart. Each topology brings in physical proximity previously characterised and new elements involved in viral replication and discontinuous transcription, therefore offering a model for facilitating distinct patterns of template switching to produce the complete SARS-CoV-2 transcriptome.

**Figure 3.**
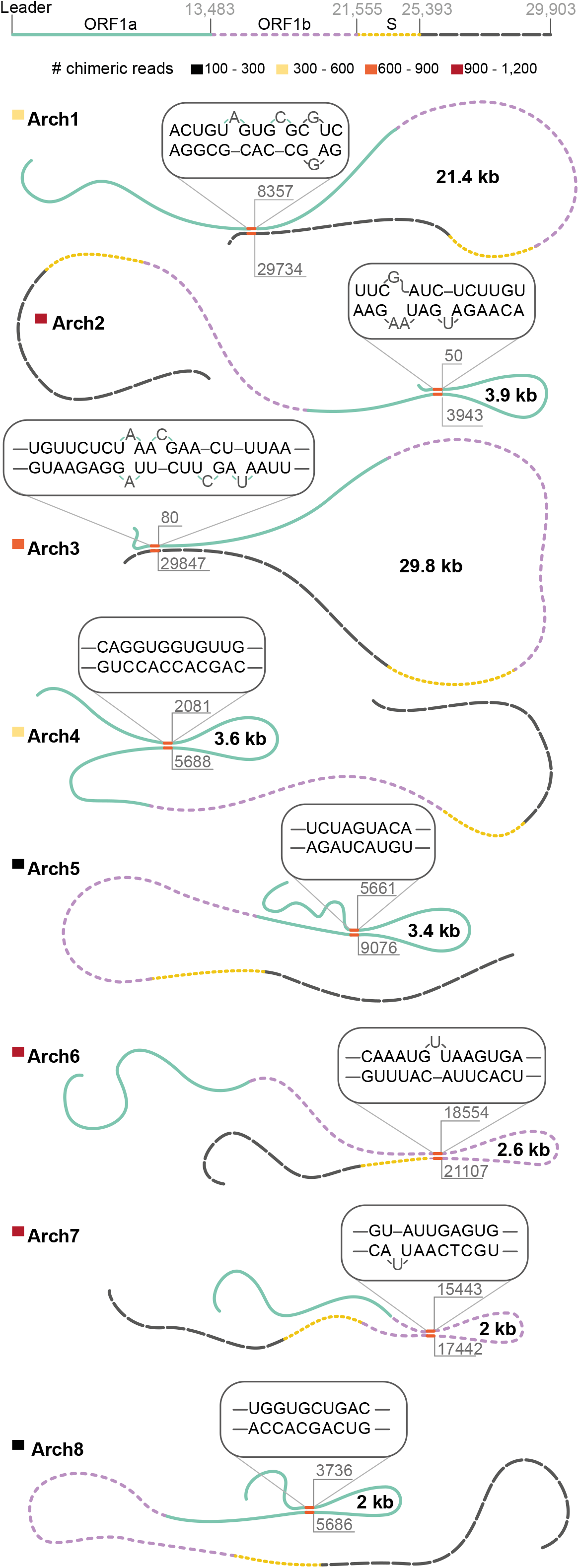
Long-range RNA-RNA interactions along the SARS-CoV-2 gRNA. RNA-RNA interactions between regions that are separated by at least 2,000 nucleotides are shown. Top panel illustrates the different patterns assigned to different parts of the genome. Coloured rectangles below the top panel and near each arch number represent the number of non-redundant chimeras supporting each conformation. The sequence of part of the base-pairing is shown above each conformation. Numbers within the loops represent the loop size. See Figure S2 and Table 1S for numbers of chimeric reads and significance of the arches.

### The SARS-CoV-2 genome engages in different interactions with cellular host RNA

The infectious life cycle of coronaviruses takes place mainly in the host cell’s cytoplasm, where many cellular RNAs reside (Sola et al., 2015). Host-virus RNA-RNA interactions regulate the replication of some RNA viruses, e.g. the interaction between hepatitis C virus and human microRNA miR-122 (Jopling et al., 2005), the interaction between Zika virus and human miR-21 (Ziv et al., 2018), and the priming of HIV-1 replication by human tRNAs (Mak and Kleiman, 1997). However, to the best of our knowledge, whether the SARS-CoV-2 gRNA or sgmRNA interact with cellular RNA is unknown. Our COMRADES method provides an opportunity to undertake an unbiased analysis of the host-virus RNA-RNA interactome (Ziv et al., 2018). We discovered site-specific interactions between the SARS-CoV-2 RNA and various cellular RNAs, especially small nuclear RNAs (snRNAs) (Figures 4A,B and S3A,B). Apart from their canonical role in splicing, snRNAs mature in the cytoplasm and may have additional biological roles (Matera et al., 2014). Along the viral gRNA, cellular snRNA interactions are mostly confined to ORF1a and ORF1b, and include site specific binding of U1, U2 and U4 snRNAs. The gRNA coding region for the sgmRNA ORFs and the UTRs are largely devoid of snRNA binding. In contrast, along the viral sgmRNA, both the N ORF and the 3′ UTR show high occupancy of U1 and U2 snRNA binding. In order to explore the conservation of these snRNA interactions in a related coronavirus, we performed COMRADES on MERS-CoV-inoculated Huh-7 cells. Similarly to SARS-CoV-2, we identified a site-specific interaction of U2 snRNA within the MERS-CoV ORF1a (Figures 4C and S3C), illustrating the evolutionary conservation of the U2 snRNA base-pairing with ORF1a of betacoronaviruses. In addition to cellular small RNAs we also detected long cellular RNAs interacting with SARS-CoV-2 RNA, although to a lesser extent. Of specific interest, the RNase MRP RNA was found to base-pair with an extended 3′ region of the sgmRNA, but not the gRNA, of SARS-CoV-2 (Figure S3D). The RNase MRP RNA has a conserved secondary structure similar to that of the RNA component of the bacterial RNase P ribonucleoprotein (RNP) complex (Dávila López et al., 2009; Welting et al., 2006). The RNase MRP RNA has a role in human pre-ribosomal RNA processing (Goldfarb and Cech, 2017), when mutated leads to a spectrum of human disease (Ridanpää et al., 2001), and has been implicated in viral RNA degradation (Jaag et al., 2011). Targeting host-virus RNA-RNA interactions provides an attractive platform for developing new antiviral therapies, as resistance would require the virus to acquire considerable mutational changes to become independent of the host RNA. However, whereas multiple tools and efforts are dedicated to identifying host-virus protein-protein interactions, the crosstalk between host and virus RNA remains largely unexplored. Coupled with the recent advancement in techniques to target RNA *in vivo*, COMRADES’s capacity to map the host-virus RNA-RNA interactome opens up new opportunities to control emerging RNA viruses. The data we present here could be valuable for the development of new targets for antiviral drugs.

**Figure 4.**
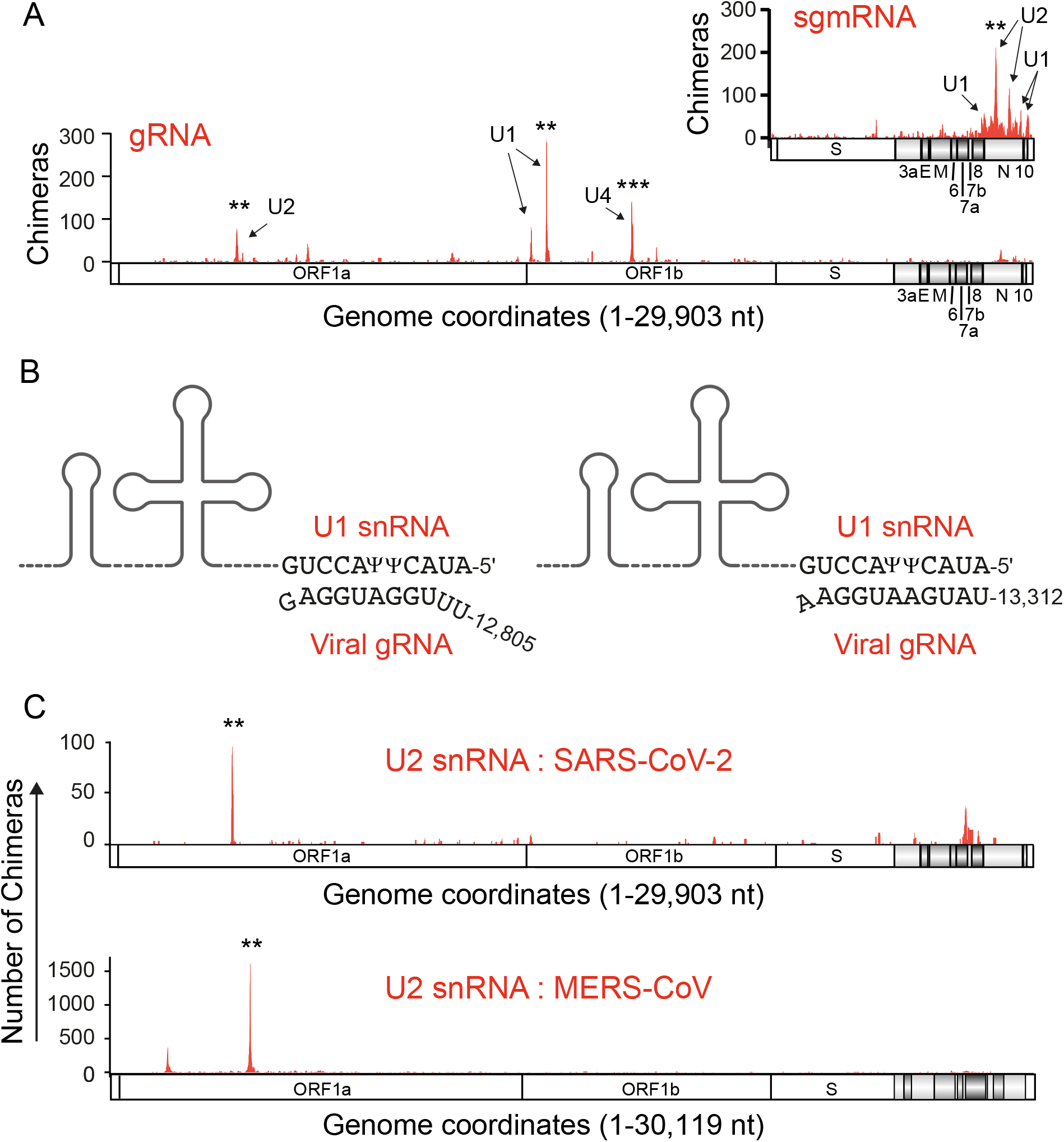
Interactions between cellular snRNAs and viral RNA. (A) snRNAs binding positions along the SARS-CoV-2 gRNA (bottom), and sgmRNA (top right). Arrows mark the binding positions of individual snRNAs. (B) Base-pairing model for the viral gRNA - U1 snRNA interaction. □ denotes pseudoUridine. (C) U2 snRNA binding positions along the SARS-CoV-2 gRNA (top), and MERS-CoV (bottom). T test p values: ** <0.05; *** <0.001. See Figure S3 for base-pairing models and for COMRADES controls.

### The UTRs of SARS-CoV-2 interact with distal genomic regions and with each other

The 5′ UTR of coronaviruses contain five evolutionary conserved stem-loop structures (denoted SL1-SL5) that are essential for genome replication and discontinuous transcription (Madhugiri et al., 2016). The 3′ UTR contains 3 structural elements important for replication: an evolutionary conserved <u>b</u>ulged <u>s</u>tem-loop (BSL) (Hsue and Masters, 1997), a partially overlapping hairpin-type pseudoknot (Goebel et al., 2004; Williams et al., 1999), and a 3′ terminal multiple stem-loop structure containing a hyper-variable region (HVR), which folds back to create a triple helix junction (Liu et al., 2013). Our analysis identified seven of these eight cis-acting elements within the UTRs (Figures 5A and S4A). However, our data did not support the folding of the stem-loop pseudoknot at the 3′ UTR. Of note, two recent studies using SHAPE methods to map the structure of SARS-CoV-2 inside cells similarly failed to identify this pseudoknot (Huston et al., 2020; Sun et al., 2020), and a previous study demonstrated the instability of this pseudoknot in the related mouse hepatitis virus (MHV) (Stammler et al., 2011).

**Figure 5.**
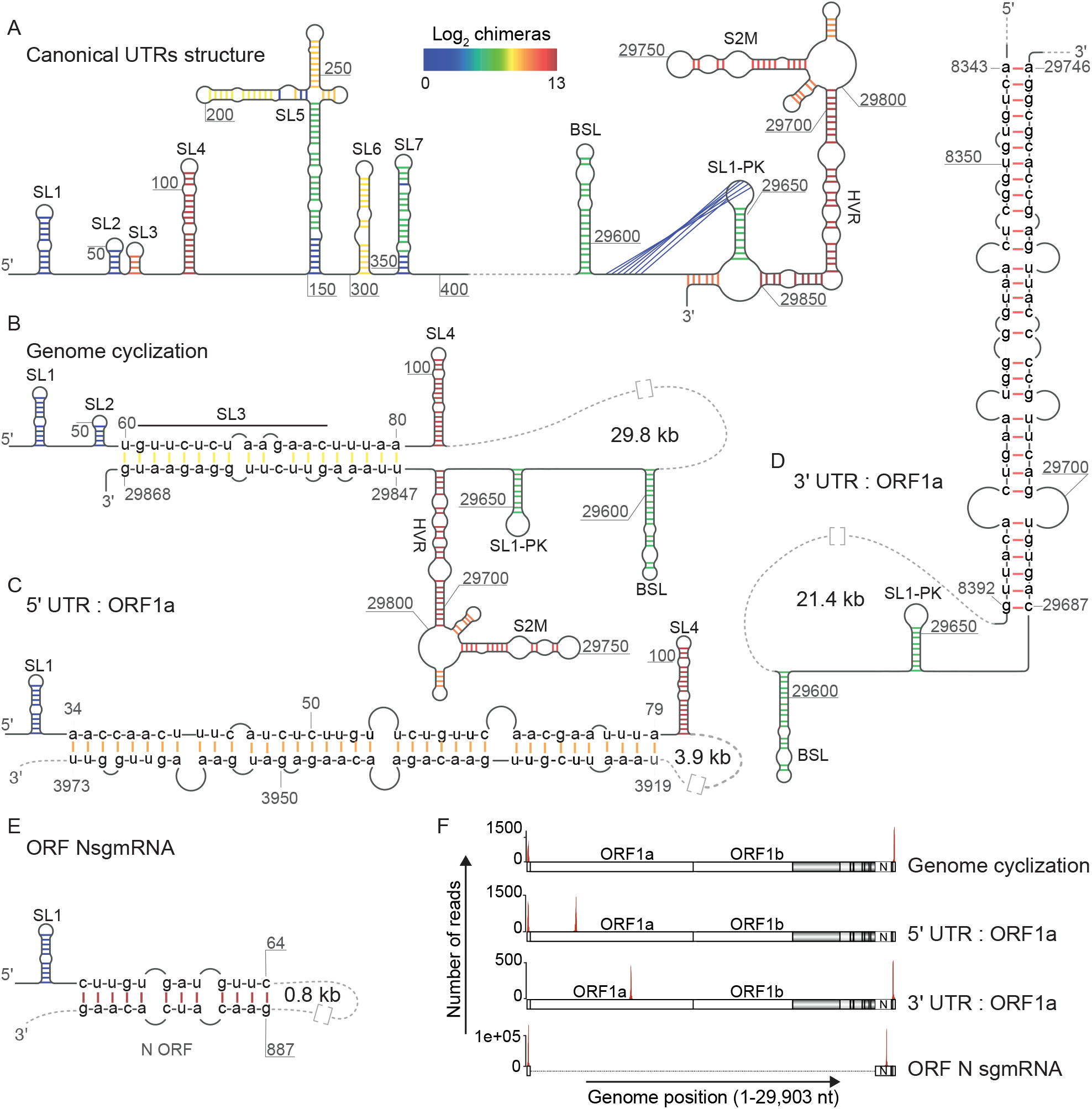
The UTRs of SARS-CoV-2 adopt alternative conformations inside cells. (A) The canonical SARS-CoV-2 UTRs structure identified in this study. Colours represent the number of non-redundant chimeric reads supporting each base-pair. (B) RNA structure corresponding to genome cyclization of SARS-CoV-2 inside cells. Colour code as in (A). (C,D) Long-distance interactions between the 5′ UTR (C) or the 3′ UTR (D) and ORF1a. Colour code as in (A). (E). Interaction between the 5′ Leader sequence and a downstream region in ORF N sgmRNA. Colour code as in (A). (F) Representation of the left- and right-side of the chimeric reads supporting the long-range interactions shown in (B-E). Numbers within loops in (B-E) represent the loops sizes. Grey arches adjacent to nucleotide sequences in (B-E) mark unpaired bases. Full sequences are available in Figure S4.

In addition to the canonical UTR structures, we provide here a direct *in vivo* evidence for genome cyclization in SARS-CoV-2, mediated by long-range base-pairing between the 5′ and 3′ UTRs (Figures 5B and S4B). This base-pairing spans a distance of 29.7 kilobases and is among the longest distance RNA-RNA interactions ever reported. Genome cyclization was previously hypothesised from mutational analyses of murine coronavirus (MHV) and was suggested to facilitate discontinuous transcription (Li et al., 2008). However, while MHV genome cyclization involves the 5′ SL1 structure, we find that in SARS-CoV-2, this process is mediated by the 5′ SL3 instead, and results in complete opening of SL3 and disruption of the triple helix junction in the 3′ UTR (Figure 5B). In agreement with this observation, SL3 of related betacoronaviruses was suggested to be weakly folded or unfolded (Chen and Olsthoorn, 2010; Li et al., 2008). Genome cyclization plays an essential role in the replication of a number of RNA viruses, including flaviviruses (Hahn et al., 1987; Ziv et al., 2018). The evolutionary selection of such a mechanism might stem from in-cell competition between intact and defective viral genomes, as it ensures that only genomes bearing two intact UTRs engage with the replication machinery. The SARS-CoV-2 genome cyclization we report here results in a complete opening of the 5′ SL3 where the <u>T</u>ranscription <u>R</u>egulating <u>S</u>equence-<u>L</u>eader (TRS-L) resides, raising the possibility that genome cyclization regulates SARS-CoV-2 discontinuous transcription, as was previously suggested for MHV (Li et al., 2008). It remains to be seen whether this base-pairing can be targeted to inhibit viral replication *in vivo*.

In addition to genome cyclization, we identified two alternative conformations involving long-distance RNA-RNA interactions between each UTR and ORF1a. These long-distance conformations result in unfolding of SL2 and SL3 in the 5′ UTR (Figures 5C and S4C), and unfolding of the terminal stem-loop in the 3′ UTR (Figures 5D and S4D). Of note, unlike the gRNA, the leader sequence within the 5′ UTR of ORF N sgmRNA is held in a single long-range conformation through base-pairing with a region 0.8 kb downstream (Figures 5E, S2B and S4E). All of the long-range interactions described above are strongly supported through chimeric reads (Figures 5F and S2A). Overall, our data demonstrate the existence of alternative, mutually exclusive UTR conformations inside cells, involving interactions between functional UTR elements and distal regions within the ORFs. We further show that the N ORF sgmRNA folds differently than the viral genome. The long-distance RNA structure map for SARS-CoV-2 provides a practical starting point to dissect the regulation of discontinuous transcription, as it identifies cis-acting elements that interact with each other to create genome topologies that favour the synthesis of the ensemble of sgmRNAs.

### A long-range structural arch encircling the SARS-CoV-2 frameshifting element (FSE-arch) is under purifying selection

RNA viruses evolve sophisticated mechanisms to enhance the functional capacity of their size-restricted genomes and to regulate the expression levels of their replicase components. In coronaviruses, one such mechanism is programmed −1 ribosomal frameshifting to facilitate translation of ORF1b which contains the viral RdRp activity, and to set a defined ratio of ORF1a and ORF1b products (Plant et al., 2010). This is mediated by a ∼120 nucleotide long cis-acting frame<u>s</u>hifting <u>e</u>lement (FSE) composed of a stem-loop attenuator, and a slippery sequence followed by a single-stranded spacer and an RNA pseudoknot (Kelly et al., 2020). It has been suggested that pausing the progression of the ribosome upstream of the pseudoknot facilitates a tandem-slippage of the peptidyl-tRNA and aminoacyl-tRNA to the −1 reading frame, thus allowing continuous translation through the stop codon at the end of ORF1a (Brierley et al., 1989). Altering the frameshifting mechanism had a deleterious effect on SARS-CoV replication (Plant et al., 2013), making the FSE an attractive target for antiviral therapy. Understanding the surrounding RNA structure and function is therefore of great importance as it might aid the design of drugs targeting the FSE. Unexpectedly, we find that the FSE of SARS-CoV-2 is embedded within a much larger, ∼1.5 kb long higher-order structure that bridges the 3′ end of ORF1a with the 5′ region of ORF1b, which we termed the FSE-arch (Figure 6A,B). To the best of our knowledge, this is the first time such a long-range structural bridge has been reported for any coronavirus, and importantly this structure is supported by the largest number of chimeric reads in our data (more than tens of thousands of non-redundant chimeric reads) (Figures S6A,B), reflecting its high folding stability *in vivo*. The FSE-arch results in a stem-loop structure encompassing 1,475 nucleotides, and bearing the FSE within it (Figures 6B and S5C). We hypothesized that if an RNA-RNA interaction is functionally important, there should be purifying selection and hence reduced nucleotide evolution rate in this region. Therefore we used a recent dataset (Firth, 2020) to explore the nucleotide conservation of the FSE-arch (Figure 6C). Strikingly, the FSE-arch is under a strong purifying selection and is among the most conserved regions within the SARS-CoV-2 genome. Consistent with this, analysing the phylogeny of the SARS-related coronavirus subgenus (taxid: 2509511) revealed two positions of covariance that support the conservation of the FSE-arch (Figure 6B). To further explore this structure experimentally, we analysed its existence in MERS-CoV. MERS-CoV shares only ∼50% sequence identity with SARS-CoV-2 (Chen et al., 2020; Lu et al., 2020), yet even so, performing COMRADES on MERS-CoV-inoculated Huh-7 cells revealed a strong evidence for an homologous FSE-arch surrounding the MERS-CoV FSE, bridging ORF1a with ORF1b (Figure 6D,E). While the mechanism governing the FSE-arch formation will require further investigation, similar long-distance interactions around the frameshifting elements of several plant RNA viruses were previously demonstrated to regulate frameshifting, possibly by assisting in back-stepping of ribosomes at the slippery sequence, and by stabilising the FSE, allowing it to refold after the passage of each ribosome (Barry and Miller, 2002; Cimino et al., 2011; Gao and Simon, 2016; Tajima et al., 2011).

**Figure 6.**
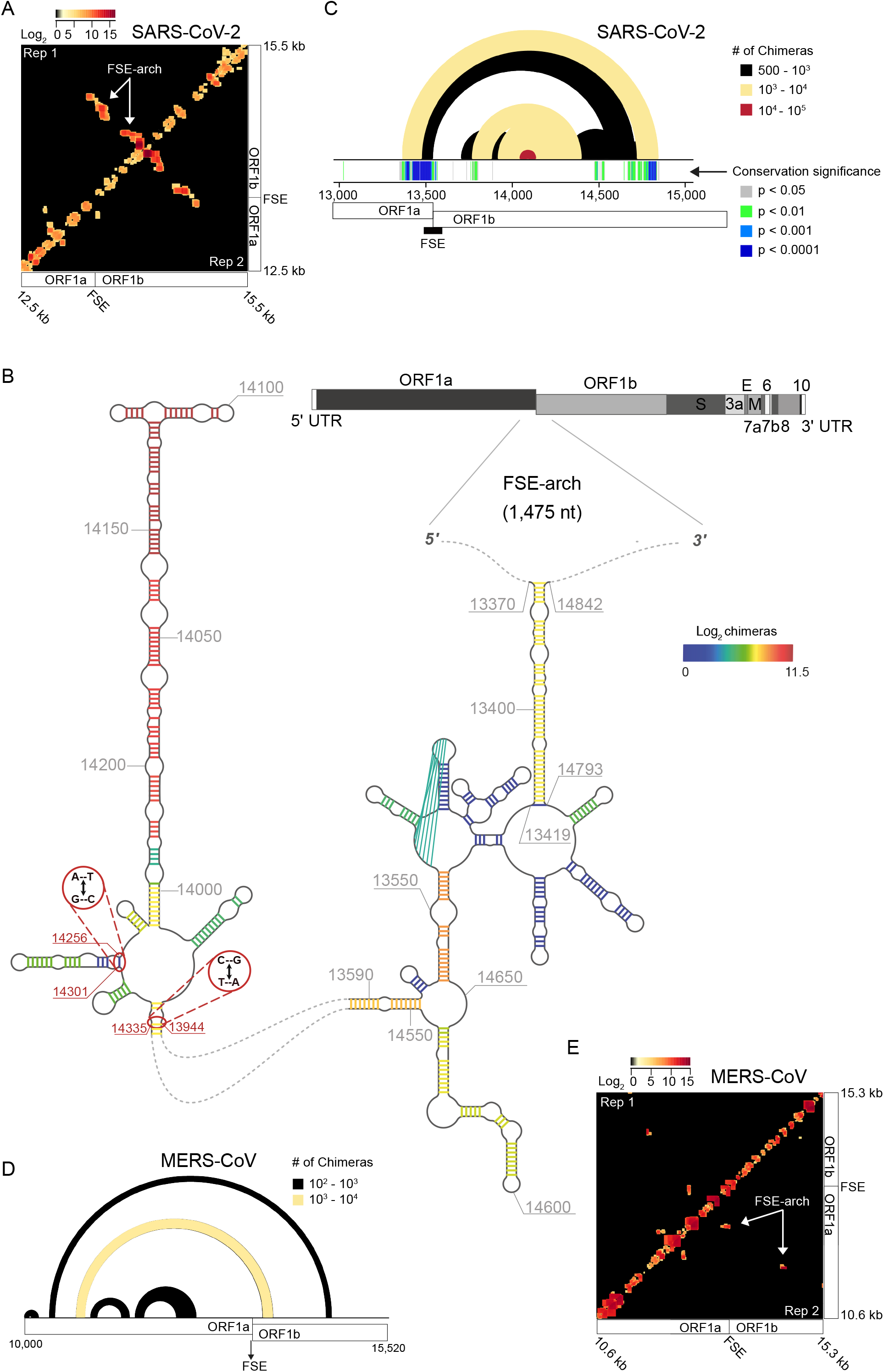
The structure of the ribosomal frame<u>s</u>hifting <u>e</u>lement <u>arch</u> (FSE-arch) inside cells. (A) Heatmap of RNA-RNA interactions around the SARS-CoV-2 FSE. Signal represents base-pairing between the genomic coordinates on the *x* and *y* axes. Top left and bottom right represent two independent biological replicates. Colour code corresponds to the number of non-redundant chimeric reads supporting each interaction. Arrows indicate the FSE-arch signal. (B) Arch plot representation of long-range RNA-RNA interactions around SARS-CoV-2 FSE. Conservation significance is shown below the arches. Black rectangle indicates the FSE position. Top Colour code corresponds to the number of non-redundant chimeric reads supporting each arch. Bottom colour code corresponds to the conservation significance (Synplot p values). (C) The structure around the SARS-CoV-2 FSE. Colours represent the number of chimeric reads supporting each base-pair. Red circles (positions 13,944 and 14,256) indicate covariation positions. (D) Arch plot representation of long-range RNA-RNA interactions around the MERS-CoV FSE. Colour code corresponds to the number of non-redundant chimeric reads supporting each arch. (E) Heatmap of RNA-RNA interactions around the MERS-CoV FSE. Signal represents base-pairing between the genomic coordinates on the *x* and *y* axes. Top left and bottom right represent two independent biological replicates. Colour code corresponds to the number of non-redundant chimeric reads supporting each interaction. Arrows indicate the MERS-CoV FSE-arch signal. See Figure 5S for statistical significance and the full sequence of the FSE-arch.

In addition to their coding capacity, nucleic acids have evolved structural capabilities to sense metabolites (Mandal and Breaker, 2004), catalyse reactions (Pyle, 1993), and interact with other cellular components. When brought in physical proximity, different combinations of cis-acting sequences can lead to new biological activities. For example, interactions between promoters and enhancers dictate the rate of transcription along the eukaryotic genome (Rowley and Corces, 2018). Great effort is being made to reveal the structural landscapes of the SARS-CoV-2 genome (Andrews et al., 2020; Huston et al., 2020; Kelly et al., 2020; Lan et al., 2020; Manfredonia et al., 2020; Ryder, 2020; Sanders et al., 2020; Sun et al., 2020). However, without deciphering the long-range connectivity, our understanding is far from being complete. Here we reveal how cis-acting elements along the coronavirus genome are folded and alternate between different topologies to create spatial combinations of functional RNA elements. The combinatorial nature of the coronavirus genome inside the host cell as discussed here, provides molecular insights into the replication, discontinuous transcription and ribosomal frameshifting machineries of coronaviruses and will facilitate the discovery of new functional cis-acting elements and the design of RNA-based antiviral therapies for SARS-CoV-2.

## Acknowledgements

The authors thank Christian Drosten and John Ziebuhrfor for providing the SARS-CoV-2 and MERS-CoV strains used in this study. We thank B. Luisi, G. Evan, and the Department of Biochemistry, University of Cambridge for facilitating the study; A. Yerra, Wellcome Trust/Cancer Research UK Gurdon Institute for critical reading and editing of the manuscript; D. Jordan for advice on statistics; I. Brierley, Department of Pathology, University of Cambridge for advising on coronavirus FSE; A.E. Firth, Department of Pathology, University of Cambridge for advising on structure conservation; S. Moss, Wellcome Trust/Cancer Research UK Gurdon Institute for assisting with risk assessments; P. Coupland and K. Kania, Genomics Core Facility, Cancer Research UK Cambridge Institute for processing the sequencing libraries; and C. Flandoli and I Kaminski for illustrations. This work was supported by Cancer Research UK grants (C13474/A18583, C6946/A14492) to E.A.M.; Wellcome grants (104640/Z/14/Z, 092096/Z/10/Z) to E.A.M.; Deutsche Forschungsgemeinschaft (DFG, German Research Foundation) – Projektnummer 197785619 – SFB 1021 grant to F.W.; the RAPID consortium of the Bundesministerium für Bildung und Forschung (BMBF) grant (01KI1723E) to F.W., and the MAD-CoV-2 consortium that has received funding from the European Union’s Horizon 2020 research and innovation program under grant agreement no. 101005026. to F.W.

## Author contributions

O.Z., L.S., and F.W. designed the study; O.Z., F.W., I.G., and E.A.M. supervised the study; L.S. carried viral infections and RNA crosslinking; O.Z. performed the COMRADES method; J.L.P. designed the analysis pipeline and analysed the data with input from O.Z. and T.K.; T.K. carried covariation analysis; O.Z., J.L.P. and E.A.M. wrote the manuscript with input from all authors.

## Material & Methods

### Cell lines

Chlorocebus sabaeus (Green monkey) VeroE6 (female, RRID:CVCL_YQ49) were purchased from American Type Culture Collection (ATCC, id: ATCC CRL-1586). VeroE6/TMPRSS2 cells (female, doi: 10.1073/pnas.2002589117) were kindly provided by Prof. Dr. Stefan Pöhlmann (German Primate Center, Göttingen). Vero E6 and VeroE6/TMPRSS2 cells were cultured in Dulbecco’s modified Eagles medium (DMEM) supplemented with 10% fetal bovine serum at 37 °C in a humidified CO2 incubator. HuH7 (Homo sapiens, male, adult hepatocellular carcinoma, RRID:CVCL_0336) cells were purchased from the Japanese Collection of Research Bioresources (JCRB) Cell Bank (JCRB No. JCRB0403) and cultured in Dulbecco’s modified Eagles medium (DMEM) supplemented with 10% fetal bovine serum at 37 °C in a humidified CO2 incubator. All cell lines were regularly examined to exclude mycoplasma contamination.

### Viral Infection and psoralen crosslinking

Infection experiments were performed under biosafety level 3 conditions. Independent biological replicates were performed using 90-120 million cells each. Titration of virus stocks was conducted using Vero E6 cells. For SARS-CoV-2 infection, VeroE6/TMPRSS2 cells (PMID: 32165541) were inoculated with SARS-CoV-2 strain München-1.2/2020/984 at MOI=2 pfu/cell for 20 hours. For MERS infection, HuH7 cells were inoculated with MERS-CoV strain EMC/2012 (GenBank: JX869059.2; PMID: 23170002) at MOI=2 pfu/cell for 20 hours. Following inoculation, cells were washed 3 times with PBS and were incubated for 20 minutes with 0.4 mg/ml Psoralen-triethylene glycol azide (psoralen-TEG azide, Berry & Associates) diluted in PBS and supplemented with OptiMEM I with no phenol-red (Gibco). Cells were subsequently irradiated on ice with 50 KJ/m2 365 nm UVA using a CL-1000 crosslinker (UVP). Cell lysis was performed by RNeasy lysis buffer (Qiagen) supplemented with DTT. Proteins were degraded using proteinase K (NEB) and RNA was purified using RNeasy maxi kit (Qiagen).

### Viral RNA enrichment

Total cellular RNA was mixed with a tiling array of 50 biotinylated DNA probes, 20 nucleotides-long each (IDT), antisense to ORF1a and ORF1b of the viral genomic RNA, and was maintained at 37 °C for 12 hours rotating in 500 mM NaCl, 0.7% SDS, 33 mM Tris-Cl pH 7, 0.7 mM EDTA, 10% Formamide. Dynabeads MyOne Streptavidin C1 (Invitrogen) were added during the final incubation hour. Beads containing the gRNA were captured on a magnet, while gRNA depleted supernatants were used for isolating the viral sgmRNA using a second tiling array of 50 biotinylated probes antisense to the sgmRNA ORFs as described in the text. Beads were washed 5 times with 2x SSC buffer supplemented with 0.5% SDS. RNA was released from the beads using 0.1 units/µl Turbo DNase (Invitrogen) at 37 °C for 30 minutes and was cleaned using RNA Clean & Concentrator (Zymo Research).

### Cross-linked RNA enrichment

Viral enriched gRNA and sgmRNA fractions were fragmented by incubating at 37 °C for 20 minutes with 0.1 units/µl RNase III (Ambion). Reactions were terminated by cleaning with RNA Clean & Concentrator (Zymo Research). Biotin was attached to cross-linked RNA duplexes by incubating with 150 µM Click-IT Biotin sDIBO Alkyne (Life technologies) under constant agitation at 37 °C for 1.5 hours. Residual Biotin sDIBO Alkyne was removed by RNA Clean & Concentrator (Zymo Research). Biotinylated RNA duplexes were enriched using Dynabeads MyOne Streptavidin C1 (Invitrogen) at the following reaction conditions: 100 mM Tris-Cl pH 7.5, 10 mM EDTA, 1 M NaCl, 0.1% Tween-20, 0.5 unit/µl Superase-In (Invitrogen). Beads were washed 5 times on a magnet with 100 mM Tris-HCl pH 7.5, 10 mM EDTA, 3.5 M NaCl, 0.1% Tween-20. RNA was eluted by adding 95% Formamide, 10 mM EDTA solution preheated to 65oC and purified using RNA Clean & Concentrator (Zymo Research).

### Proximity ligation and crosslink reversal

Each RNA sample was divided in two: one half was used for proximity ligation and then crosslink reversal (i.e. COMRADES sample), while in the other half, crosslink reversal was done before proximity ligation (i.e. control sample). Prior to proximity ligation, RNA was denatured by briefly heating to 95 °C. Proximity ligation was done under the following conditions: 1 unit/µl RNA ligase 1 (New England Biolabs), 1x RNA ligase buffer, 50 µM ATP, 1 unit/µl Superase-in (Invitrogen), final volume: 200 µl. Reactions were incubated for 16 hours at 16 °C and were terminated by cleaning with RNA Clean & Concentrator (Zymo Research). Crosslink reversal was done by irradiating the RNA on ice with 2.5 KJ/m2 254 nm UVC using a CL-1000 crosslinker (UVP).

### Sequencing library preparation

Library preparation was done as described in (Ziv et al., 2018), using 6N unique molecular identifiers to eliminate PCR biases. Pre-adenylated adapters were used and all ligation reactions were carried without ATP to reduce ligation artefacts. All libraries and controls went through 13 PCR cycles using KAPA HiFi HotStart Ready Mix (KAPA Biosystems). PCR products were size-selected on a 1.8% agarose gel before loading on a Novaseq (Illumina) for paired-end 150 bp runs. Total of ∼1.6 billion sequences were achieved for this study.

### Data Pre-processing

Data pre-processing was performed according to (Ziv et al., 2018). In brief, raw paired-end reads were trimmed for adaptors and checked for quality using cutadapt (Martin, 2011). Trimmed paired-end reads were assembled into single reads using the program pear (https://cme.h-its.org/exelixis/web/software/pear/doc.html). PCR duplicates were removed using unique molecular identifiers via collapse.py (https://gitlab.com/tdido/tstk). Chimeric reads were identified and annotated to the respective genome using hyb (Travis et al., 2014). SARS-CoV2 samples were processed using the Chlorocebus sabaeus reference genome (ChlSab1.1) with the addition of the SARS-CoV-2 sequence (NC_045512.2). MERS samples were processed using the Homo sapiens reference genome (GRCh38) with the addition of MERS (NC_019843.3).

### Clustering of chimeras into chimeric groups

Due to crosslinking and fragmentation, the COMRADES data can provide redundant structural information whereby the same *in vivo* structure produces sequencing reads differing by a few nucleotides. This results in increased computation load of folding each chimeric read separately. To overcome this issue, and to gain better structure predictions, the reads were clustered into chimeric groups. Each chimeric read is composed of a left side (*L*) and right side (*R*), each originated from a different position along the gRNA or sgmRNA. Each chimeric read can therefore be described as (*g*): the genomic distance between *L* and *R*, and chimeric reads that originated from the same structure will have a similar *g* and can be clustered based on their *g* values.

Clustering of chimeric reads that originated from the same structure was performed using a network-based approach whereby an adjacency matrix is created for all chimeric reads based on the nucleotide difference between their *g* values (*Deltagap*).

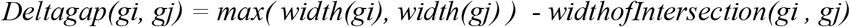

This results in *Deltagap=0 for* identically overlapping gaps, and increasing *Deltagap values for* chimeric reads that share less overlapping sites. The clustering was performed twice per sample, once for the chimeric reads that represent short stem structures (*g* <= 10 nt) and once for chimeric reads that represent long distance interactions (g > 10 nt). Short range interactions weights were calculated as:

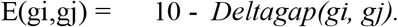

This allows exactly overlapping gaps to have the highest weight, and gaps with no overlaps to have a weight of 0. For long range interactions weights were calculated as:

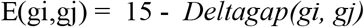

Long range interactions with weights lower than 0 were set to 0, meaning that gaps that differ by more than 15 nucleotides could not be considered as part of the same chimeric group. The weighted graphs created for long- and short-range interactions consisted of *g* as vertices and weights as edges:

G = (V,E) using iGraph (http://www.interjournal.org/manuscript_abstract.php?361100992). To identify densely connected subgraphs (communities) with chimeric groups containing chimeric reads that originated from the same structure, we clustered the graph using random walks with the cluster_waltrap function (steps = 2) from the iGraph package. Chimeric groups containing less than 10 chimeric reads were discarded. Chimeric groups often contained a small amount of longer *L* or *R* sequences due to the random fragmentation in the COMRADES protocol. To avoid introducing biases in the folding results, clusters were trimmed to the region from *L* to *R* for which evidence in the cluster is higher than the mean evidence - 2 standard deviations.

### Folding

Folding of the chimeric groups was performed using the Vienna package (Lorenz et al., 2011). For short range chimeric groups RNAFold was used (with default parameters) and for long range chimeric groups RNADuplex was used (with default parameters).

### Covariation analysis

Complete Sarbecovirus sequences were taken from the NCBI Virus Database website (taxid: 2509511, 17 June 2020 version). Sequences containing unidentified nucleotides were discarded. Four sequence sets for MSA (multiple sequence analysis) were generated:

I. MSA-1-SARSrel-3515seq: includes all complete non-redundant Sarbecovirus sequences. MUSCLE ((Edgar, 2004)), (default parameters).
II. MSA-2-SARSrel-3515seq: Includes all complete non-redundant Sarbecovirus sequences containing only the four canonical nucleotide identifiers. MUSCLE (parameters, -maxiters 2).
III. MSA-3-SARSrel-137seq: pairwise distances between the sequences in *(I)* were calculated using the mbed function in the kmer package (Blackshields et al., 2010). The “seeds” attribute was used to extract the sequence indices of all seed sequences. These seed sequences were included in this multiple sequence alignment. This resulted in a smaller sequence set but with a representative variation as (I). MUSCLE (default parameters).
IV. MSA-4-SARSrel-559seq: The unaligned sequences used to generate *(II)* were divided into seven smaller sequence sets (six 500-sequences sets and one 515-sequence set). The seed sequences (“seeds” attribute of the mbed function in the kmer package (https://cran.r-project.org/package=kmer)) for all seven sets were combined in a new sequence set to be aligned to make up multiple sequence alignment *(IV)*. This resulted in a sequence set with less sequences than (I), but more than (III) and representative variation as (I). MUSCLE (default parameters).

In all cases the NCBI reference genome for SARS-CoV-2 (NC_045512.2) was used as reference. For each of the four sequence alignments, the following steps were taken:

I. SARS-CoV-2 COMRADES cluster coordinates. For long-range clusters (defined above) the segments defined by the coordinates of the left and right side of the respective cluster were extracted from the MSA, and fused together to form a smaller MSA containing only the aligned left and right side sequences. For short-range clusters (defined as above) the whole region defined by the start position of the left side and the end position of the right side was extracted. Those segments of the full MSA will be referred to as “cluster alignments”. In both cases, any sequence starting with more than 10 empty positions was removed from that cluster alignment.
II. The cluster alignments were analyzed with R-Scape (Rivas et al., 2017) using default parameters.
III. The R-Scape output for each candidate co-varying pair includes an E-value statistical score (probability of a false positive result for the respective position pair in the cluster alignment). The default significance level of 0.05 was kept, so only position pairs with E-values smaller than 0.05 were considered in the subsequent analysis.
IV. Results tables of R-scape output combined with the corresponding coordinates in the non-aligned SARS-CoV-2 reference sequence, as well as nucleotide combination frequencies at the two positions across the alignment. We defined secondary base-pairing frequency as the percentage of sequences in which the pair of nucleotides differed from the most common base-pair type at the position but could still form a base-pair. For example, an imaginary pair of nucleotides has the composition:

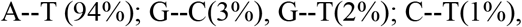

And its secondary base-pairing frequency is 83.3%. For further steps of the analysis (folding predictions), we consider only candidate pairs with ≥90% secondary base-pairing frequency.

### Covariation analysis of sgmRNA chimera clusters

The alignments described above were shortened to include the leader sequence fused to the full-length of mRNA-S, and were subsequently used here. A modified version of the code used from full genome chimera clusters was used.

### Sequence conservation analysis of the extended FSE structure

Genome conservation data analyzed with synplot2 (Firth, 2014) was taken from (Firth, 2020). These data were aligned with our structural data and displayed in Figure 6C.

