## Supplemental Figure 1 for "The short- and long-range RNA-RNA Interactome of SARS-CoV-2"

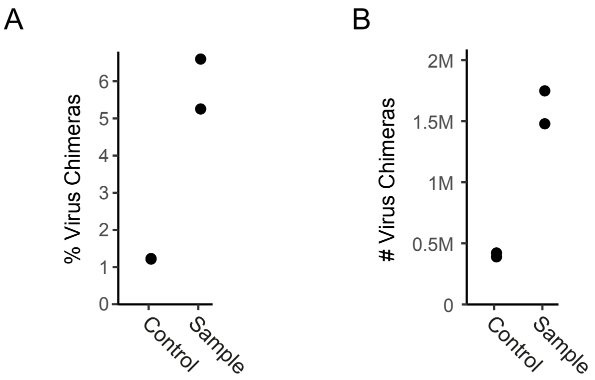


**Supplementary Figure 1, related to Figure 1. The COMRADES method.**
(A) Percentage of chimeric reads in the samples and control experiments.
(B) Number of chimeric reads in the samples and control experiments.

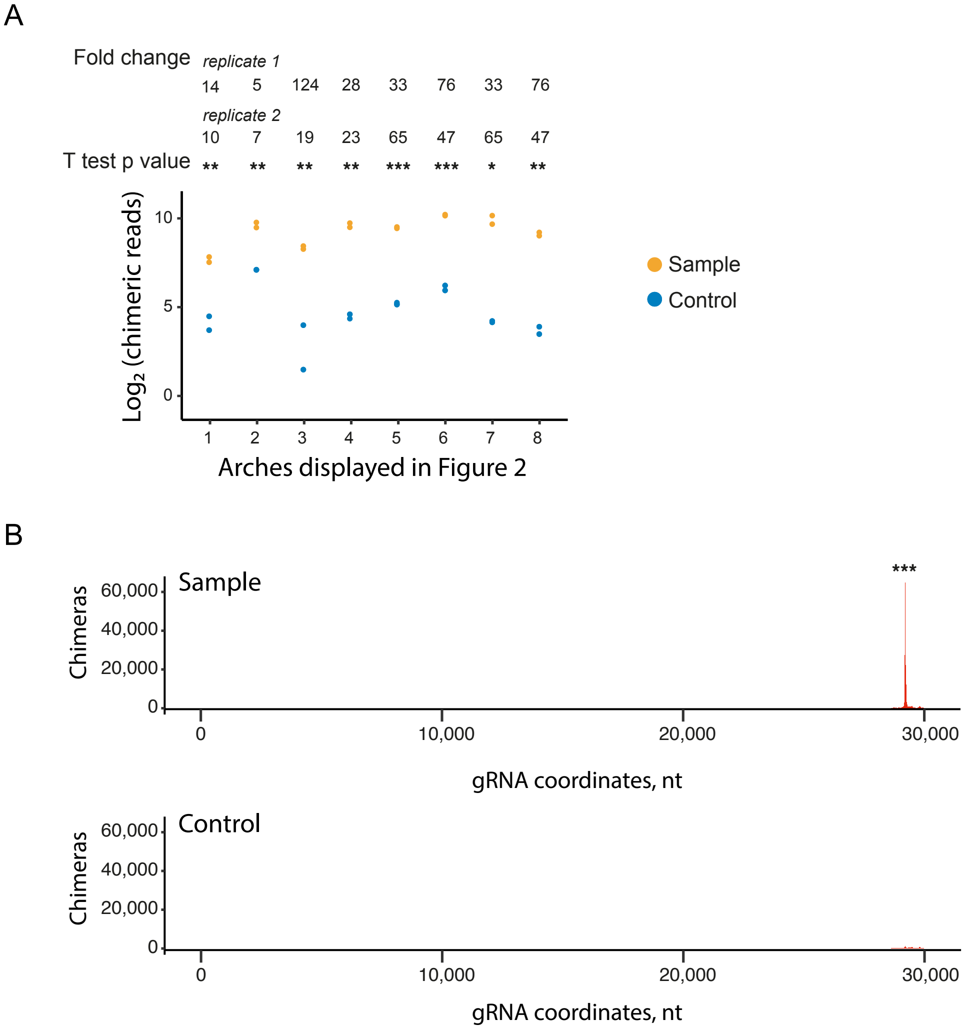


**Supplementary Figure 2, related to Figure 3. Long-range RNA-RNA interactions along the SARS-CoV-2 gRNA and N ORF sgmRNA**
(A) Number of chimeric reads supporting the arches shown in Figure 2 in COMRADES samples and controls libraries. $Log_2$ values are shown. Fold changes represent the ratios of chimeras in sample / chimeras in control for each biological replicate. T test p values indicate the significant of each arch (* < 0.1,  ** < 0.05,  *** < 0.01).
(B) Interactions of the leader sequence with a downstream position in ORF N sgmRNA. Number of chimeric reads supporting this base-pairing is shown. *** denotes T test p value < 0.01

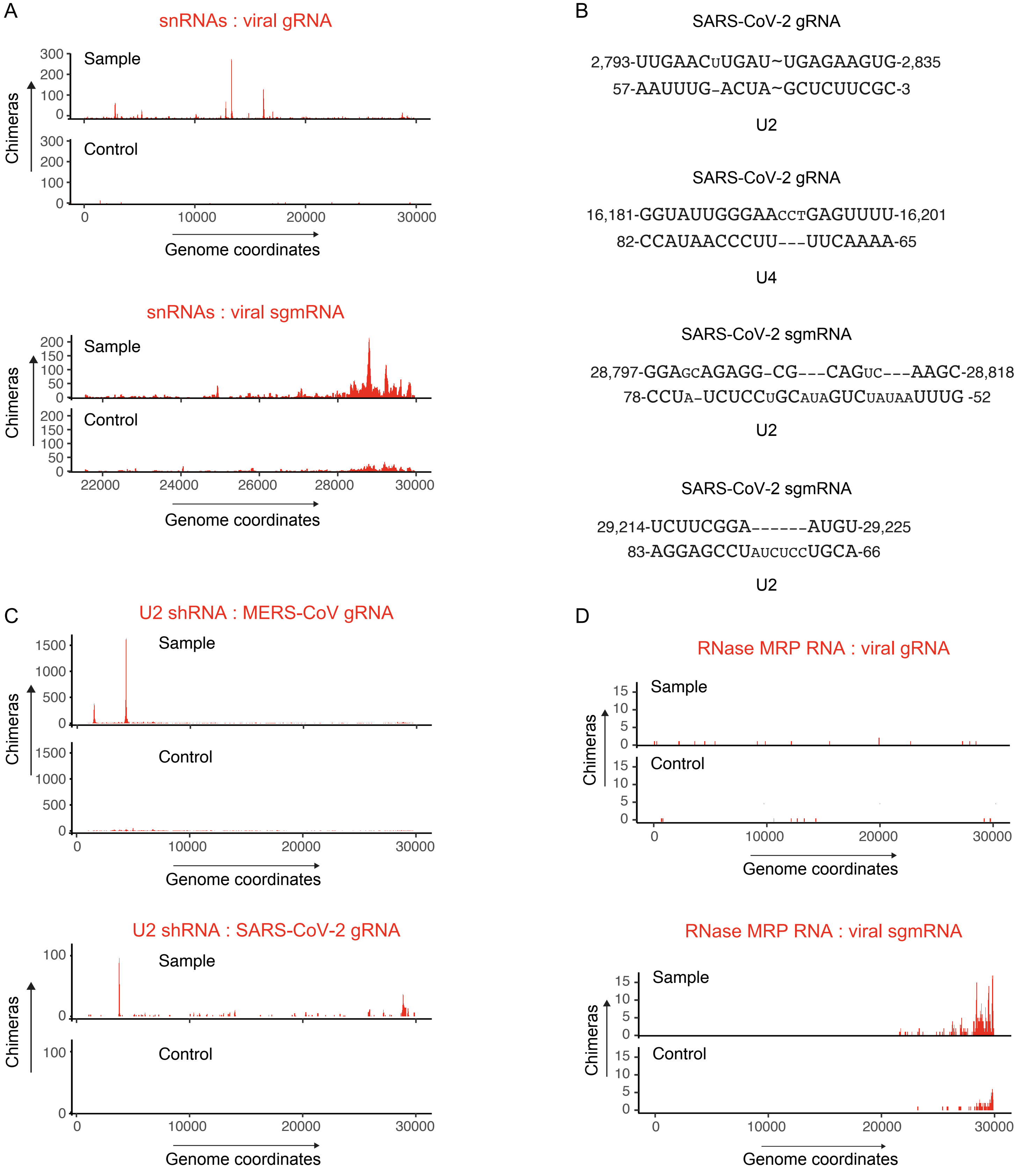


**Supplementary Figure 3, related to Figure 4. Interactions between cellular and viral RNA**
(A) snRNAs binding positions along the SARS-CoV-2 gRNA (top) and sgmRNA (bottom) and their COMRADES-controls.
(B) Base-pairing models for the interactions between viral RNA (top strand) and host snRNAs (bottom strand)
(C) U2 snRNA binding position along MERS-CoV gRNA (top) and SARS-CoV-2 gRNA (bottom) and their COMRADES controls
(D) RNase MRP binding positions along the SARS-CoV-2 gRNA (top) and sgmRNA (bottom) and their COMRADES controls.

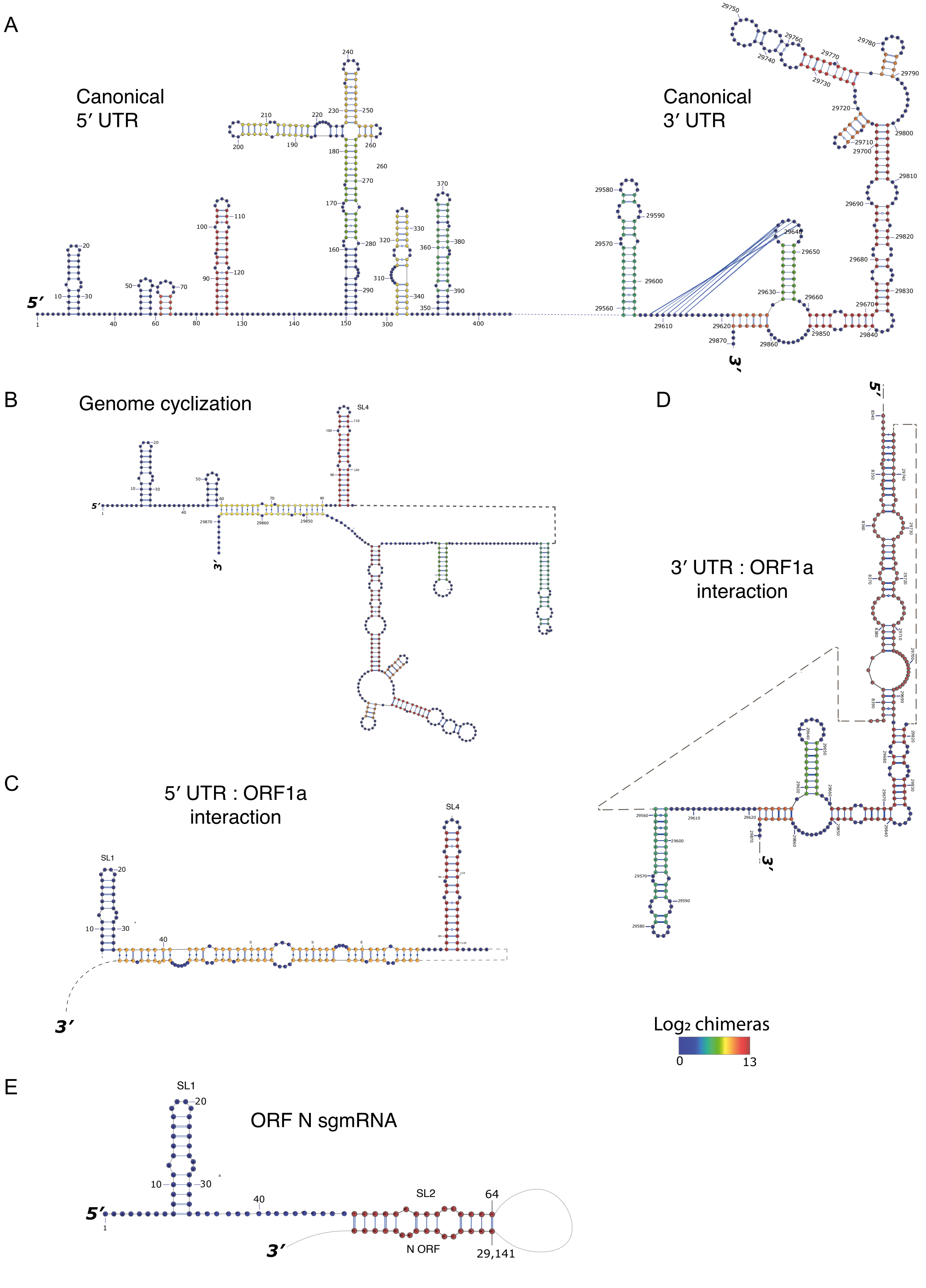


**Supplementary Figure 4, related to Figure 5. The UTRs of SARS-CoV-2 adopt alternative conformations inside cells**

(A-E) Detailed representation of the canonical UTRs structure (A); genome cyclization (B); UTRs binding to ORF1a (C,D); and ORF N sgmRNA conformation (E). Colour code represents the number of non-redundant chimeric reads supporting each base-pair.

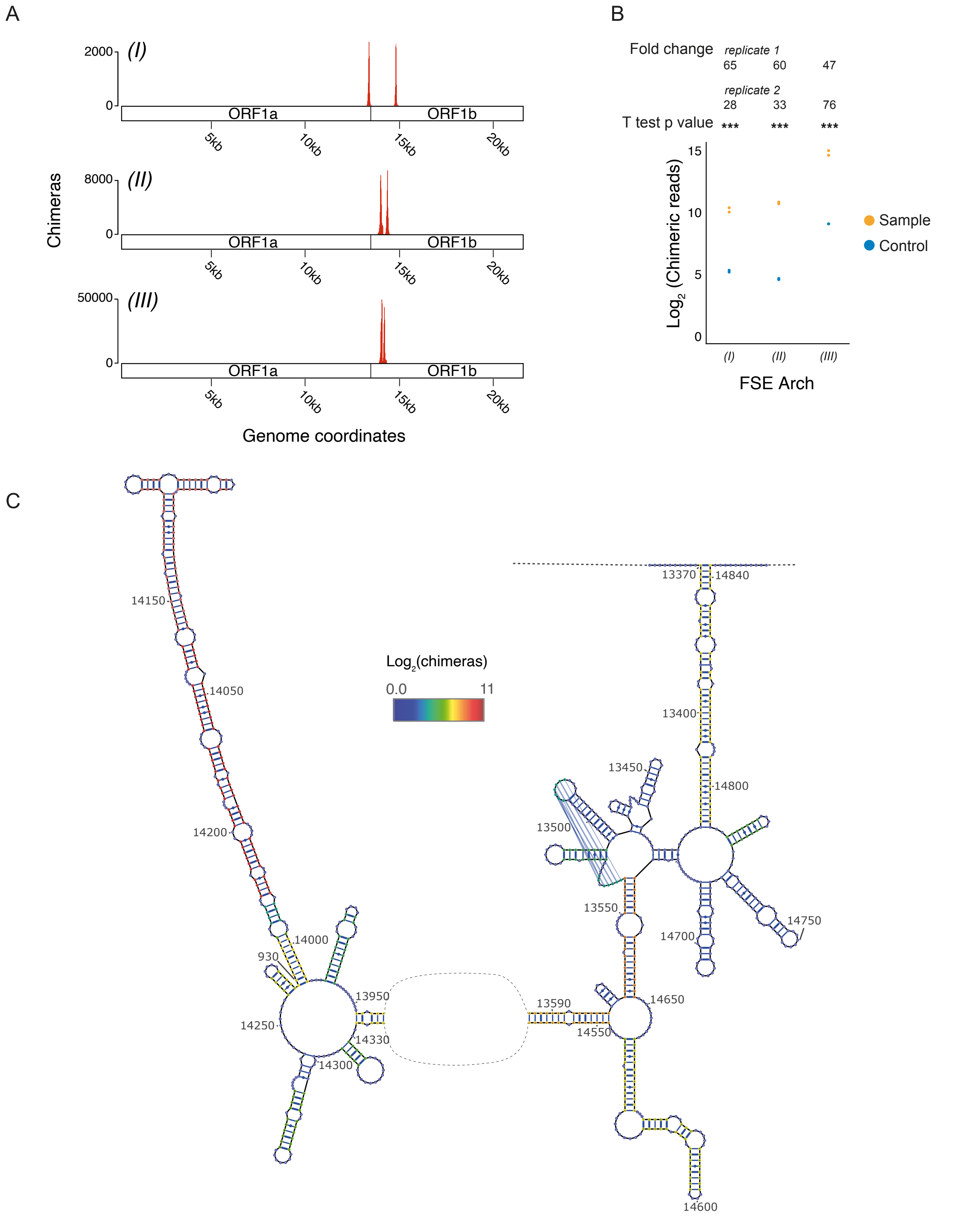


**Supplementary Figure 5, related to Figure 6. The structure of the SARS-CoV-2 ribosomal frameshifting element arch (FSE-arch) inside cells**

(A) Representation of the left- and right-side of the chimeric-reads supporting the FSE-arch. *(I)*-*(III)* correspond to the gRNA positions marked in (C).

(B) Number of chimeric reads supporting the FSE-arch in samples and controls. $Log_2$ values are shown. Fold changes represent the ratios of chimeras in sample / chimeras in control for each biological replicate. T test p values indicate the significant of each position (*** < 0.01). *(I)*-*(III)* correspond to the gRNA positions marked in (C).

(C) Detailed representation of the FSE-arch structure. Colour code represents the number of non-redundant chimeric reads supporting each base-pair.

**Supplementary Table 1. Prevalence of chimeras supporting the long-range interactions shown in Figure 3.**

| Arch name | Long-range chimeras[a] | Short-range chimeras[b] | Ratio of Long-range / Short-range chimeras |
|---|---|---|---|
| 1 | 442 | 14,506 | 3.0% |
| 2 | 1,873 | 5,980 | 31.3% |
| 3 | 704 | 5,980 | 11.8% |
| 4 | 1,690 | 13,546 | 12.5% |
| 5 | 1,529 | 9,050 | 16.9% |
| 6 | 2,493 | 7,447 | 33.5% |
| 7 | 2,102 | 16,531 | 12.7% |
| 8 | 1,193 | 11,766 | 10.1% |

[a]Long-range chimeras supporting each arch
[b]Chimeras supporting alternative, short-range interactions within the arch regions
